# Neural decoding of visual stimuli varies with fluctuations in global network efficiency

**DOI:** 10.1101/107888

**Authors:** Luca Cocchi, Yang Zhengyi, Zalesky Andrew, Stelzer Johannes, Luke Hearne, L. Gollo Leonardo, Jason B Mattingley

**Author notes:** Corresponding authors: Luca Cocchi, QIMR Berghofer Medical Research Institute. Herston, 4006, QLD, Australia.; or.

## Abstract

Functional magnetic resonance imaging (fMRI) studies have shown that neural activity fluctuates spontaneously between different states of global synchronization over a timescale of several seconds. Such fluctuations generate transient states of high and low correlation across distributed cortical areas. It has been hypothesized that such fluctuations in global efficiency might alter patterns of activity in local neuronal populations elicited by changes in incoming sensory stimuli. To test this prediction, we used a linear decoder to discriminate patterns of neural activity elicited by face and motion stimuli presented periodically while participants underwent time-resolved fMRI. As predicted, decoding was reliably higher during states of high global efficiency than during states of low efficiency, and this difference was evident across both visual and non-visual cortical regions. The results indicate that slow fluctuations in global network efficiency are associated with variations in the pattern of activity across widespread cortical regions responsible for representing distinct categories of visual stimulus. More broadly, the findings highlight the importance of understanding the impact of global fluctuations in functional connectivity on specialised, stimulus driven neural processes.

## Introduction

Recent studies have highlighted the dynamic nature of neural activity in the absence of explicit task demands (Allen et al. 2014; Breakspear 2004; Chang and Glover 2010; Karahanoglu and Van De Ville 2015; Ponce-Alvarez et al. 2015). By combining time-resolved functional magnetic resonance imaging (fMRI) and tools from the field of network science, it has recently been shown that neural activity fluctuates spontaneously in and out of global synchronization over a timescale of several seconds (Zalesky et al. 2014). Such fluctuations in hemodynamic activity generate transient states of high and low correlation across distributed cortical areas, resulting in global changes in network efficiency. Global efficiency is a metric used to quantify the capacity of a complex network to share information between spatially segregated communities (Latora and Marchiori 2001). Efficiency computed in functional brain networks provides an index of the brain’s capacity to support the parallel transfer of information (Achard and Bullmore 2007; Sporns 2011).

It has been hypothesized that transitions between high and low states of global efficiency might influence patterns of activity in local neuronal populations. Specifically, results from computational and empirical work suggest that slow fluctuations in neural activity underpinning efficiency may play a critical role in maintaining global network stability while facilitating generation of, or access to, sensory representations (de Pasquale et al. 2015; Gollo et al. 2015; Kringelbach et al. 2015; Murray et al. 2014; Shine et al. 2016). Thus, for example, neural responses associated with the presentation of a given visual input might be associated with distinct patterns of local neuronal activity under high versus low states of global network efficiency. Alternatively, ongoing slow fluctuations in global network efficiency may represent a mechanism of the brain’s energy homeostasis (Bullmore and Sporns 2012; Zalesky et al. 2014) that has no significant impact on patterns of neural activity associated with the encoding, maintenance or retrieval of sensory information. To date, however, it remains unknown whether slow fluctuations in global network efficiency are associated with changes in local activity elicited by the presentation of distinct sensory inputs, independent of explicit task demands.

To address this issue, we combined a passive visual monitoring paradigm with multiple sessions of sub-second resolution functional magnetic resonance imaging (fMRI) (**Fig. 1**). The sub-second temporal resolution of the MR protocol allowed for robust characterization of fluctuations in global network efficiency. During scanning, participants were asked to relax and keep their gaze on a central fixation cross. Participants were explicitly instructed to ignore a sparsely presented stream of face and motion stimuli that appeared at fixation (see SMovie for trial example). To determine the impact of expected slow changes in global neural efficiency on faster stimulus-driven processes, we analysed the data using approaches sensitive to global changes in dynamic connectivity (Zalesky et al. 2014) and a linear classifier allowing the isolation of patterns of neural response evoked by two distinct categories of visual stimuli (Stelzer et al. 2013). Neutral human faces and patches of visual motion were selected as stimuli because they are known to evoke specific and robust multivoxel patterns of activity in the visual system, specifically in the fusiform gyrus (Haxby et al. 2001) and medial temporal area [MT, (Hong et al. 2012)], respectively. Due to constraints imposed by the sparse, event-related design and relatively short scan duration (20 mins), we used human faces as one category, and radial-dot motion as the other, to maximize the strength and reliability of decoding across widespread regions of the cortex. We reasoned that if changes in global efficiency affect local activity patterns elicited by the face and motion stimuli, classification accuracy should be different when these visual events appear during states of high versus low network efficiency (**Fig. 1**).

**Figure 1.**
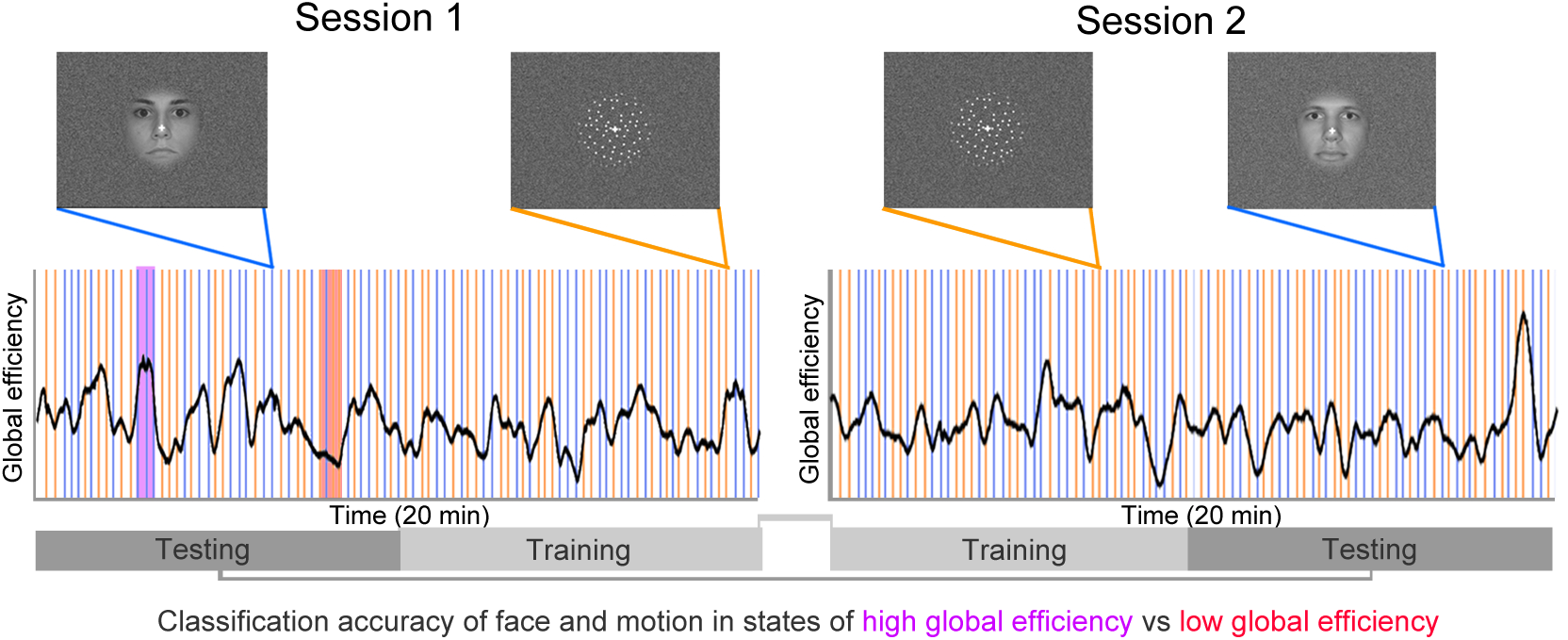
Schematic representation of the experimental design. Participants underwent two separate fMRI sessions. In each session they were asked to lie in the scanner, relax, and keep their eyes on a fixation cross presented at the centre of the screen. Participants were instructed to ignore stimuli (face and motion) presented at the centre of the visual display. Stimuli were presented sparsely (one every 12 seconds on average) and in a pseudorandom order. To test the hypothesis of altered local activity patterns during epochs of high (purple) versus low (red) global efficiency, we used a linear classifier to isolate brain regions showing significant decoding between the two stimulus categories. Trials in each fMRI session were divided in two and intermingled to train and test the classifier on independent datasets. We characterised dynamic fluctuations in global (whole-brain) efficiency (black line) using tools from network science (Zalesky et al. 2014). We tested for differences in decoding accuracy when stimuli occurred during states of high efficiency versus states of low efficiency.

We began by isolating brain areas in which mean activity (Friston et al. 1994), or multivoxel patterns of activity (Kriegeskorte et al. 2006), differentiated face and motion stimuli. Next, we used a validated, time-resolved approach to characterise spontaneous fluctuations in whole-brain functional connectivity and global network efficiency (Zalesky and Breakspear 2015; Zalesky et al. 2014). We then assessed whether the fluctuations in global neural states, characterised as high or low efficiency, were associated with differences in decoding of local neuronal activity elicited by the face and motion stimuli.

## Materials and Methods

The study was approved by The University of Queensland Human Research Ethics Committee. Written informed consent was obtained for all participants. Data from 26 healthy adult participants were acquired on a 3T Siemens Trio MR scanner equipped with a 32-channel head coil. Participants underwent two 20-minute fMRI scans, separated by an anatomical scan, all within a single 60-minute experimental session, as outlined in detail below. We performed two fMRI scans per participant so that the decoding algorithm could be trained and tested on independent datasets (**Fig. 1**).

During the two fMRI sessions, participants were instructed to maintain their gaze on a fixation cross presented in the middle of the screen, to relax, and to avoid focusing on any particular thought. Participants were informed that images of neutral human faces or radially moving dots would sporadically and gradually appear and disappear on the background of the white fixation cross, and that they should ignore them as no response was required. Participants’ eyes were constantly monitored during data collection to ensure they did not fall asleep or become excessively drowsy (Fukunaga et al. 2006; Tagliazucchi and Laufs 2014). Six participants were excluded because they closed their eyes for prolonged periods (> 5 sec) in at least one of the two fMRI sessions. All analyses were therefore performed on a final sample of 20 participants (mean 28, S. D. ± 7 years; 12 females).

### Stimulus materials

Stimuli were presented using Psychtoolbox 3 (http://psychtoolbox.org/, installed on a Dell Optiplex 960 with Windows XP) and back-projected onto a screen positioned at the head end of the scanner by a liquid crystal display (LCD) projector (Canon sx80). Twenty-four human faces displaying a neutral expression and 24 radial dot motion stimuli were presented in a sparse and pseudo-random order in each fMRI run (**SMovie**).

Stimuli were first grouped in random blocks of 3 faces and 3 motion stimuli. Next, blocks were concatenated and modified such that the same stimulus category (face or motion) could not appear more than 3 times in a row. Different face and motion stimuli were used across the two fMRI runs. The minimum inter-stimulus-interval between the stimuli was 9 seconds, and the maximum was 15 seconds (average of 12 seconds, standard deviation ± 2.8 seconds). Stimuli were presented within a circular region (approximately 8º visual angle) centered at fixation. Neutral faces were selected and modified (i.e., converted to black and white, standardized for luminance, contrast, eye position at the centre of the image, and eyes aligned to the horizontal axis of the fixation cross) from a standard dataset (Ebner 2008). Radial motion stimuli were generated using Psychtoolbox 3, and consisted of white dots (~ 0.2º diameter) moving radially outward from the centre of the screen. The velocity of each dot was proportional to its distance from the centre of the screen (6/200 pixels per frame multiplied by pixel distance from the centre). Once the dots reached the border of the 8º central region, they were re-spawned at a new random position within the region. Both stimulus types began at 0% visibility (100% transparent) and gradually increased in contrast over 1000 ms until they became fully visible (100% opaque). Stimuli remained visible for 2000 ms, before progressively reducing in contrast over 1000 ms until they disappeared (see SMovie for examples).

### Functional MR data

For both fMRI sessions, whole brain echo-planar images were acquired using a multi-band sequence (acceleration factor of four) developed and optimized in collaboration with Siemens Healthcare (Siemens WIP770 sequence). Each imaging sequence lasted approximately 20 minutes (44 axial slices, 1705 volumes, slice thickness = 3 mm; matrix size = 64 × 64, in-plane resolution = 3 × 3 mm^2^, repetition time = 700 ms, echo time = 28 ms, flip angle = 54°, FOV = 192 × 192 mm^2^). T1 images were acquired using the following parameters: 192 axial slices; slice thickness = 0.9 mm; matrix size = 512 × 512, in-plane resolution = 0.4492 × 0.4492 mm^2^, repetition time = 1900 ms, flip angle = 9°, echo time = 2.32 ms, FOV = 230 × 230 mm^2^.

### Preprocessing

Imaging data were preprocessed using the Matlab (MathWorks, USA) toolbox *Data Processing Assistant for Resting-State fMRI A* [DPARSF, (Chao-Gan and Yu-Feng 2010)]. The first 14 image volumes (9.8 seconds) were discarded to allow tissue magnetization to reach a steady-state and let participants adapt to the scanner environment. DICOM images were first converted to Nifti format. T1 images were manually re-oriented, skull-stripped, and co-registered to the Nifti functional images. Segmentation and the DARTEL algorithm were used to improve the estimation of non-neural signal (see below) and spatial normalization (Ashburner 2007). Single-subject functional images were normalized to standard MNI space and smoothed using a Gaussian function with an 8 mm full width at half maximum (FWHM) kernel. Data processing steps also involved filtering [0.01-0.15 Hz; this low-frequency component of the BOLD signal was selected because it is known to be sensitive to both rest and task-based functional connectivity (Bassett et al. 2015; Boubela et al. 2013; Niazy et al. 2011; Sun et al. 2004)], exclusion of undesired linear trends and regression of signals from the six head motion parameters from each voxel’s time series. The CompCor method (Behzadi et al. 2007) was used to regress out residual signal unrelated to neural activity such as heart rate and respiration (i.e., five principal components derived from single subject white matter and cerebrospinal fluid masks generated by the segmentation). This method has been shown to be more efficient in removing physiological noise than methods requiring external monitoring of such parameters (Behzadi et al. 2007). Note that we previously demonstrated that fluctuations in global network efficiency are not due to changes in physiological noise (Zalesky et al. 2014). To further control for potential confounds related to micro-head motion, frames with a volume-level mean of frame-to-frame displacements greater than 0.25 mm (including the preceding and the two subsequent frames) were removed and interpolated using a cubic spline function (Power et al. 2012; Power et al. 2014). Overall, the number of interpolated volumes using this stringent frame-to-frame movement threshold was small and similar between the two fMRI sessions (interpolated volumes session 1: 2.8%, session 2: 3.5%, p = 0.5).

### General linear model

Statistical analyses were performed by adopting the general linear model (GLM) as implemented in the Matlab toolbox SPM8 (http://www.fil.ion.ucl.ac.uk/spm/software/spm8/). At the first level (within-subject analysis), the conditions of interest were modeled as boxcar functions convolved with a canonical hemodynamic response function. The first-level model comprised regressors for the face epochs (4 seconds) and the motion epochs (4 seconds). The resulting first-level contrast images (motion > faces and faces > motion) were carried over to second-level random-effects analyses (one-sample t-test, p < 0.05 FDR corrected at the cluster level, with an uncorrected high threshold of p = 0.02. This lenient search threshold was used to generate a comprehensive mask of the regions of interest for the classifier – see below for details).

### Time-resolved analysis of global functional connectivity

Time-resolved functional connectivity was estimated exactly as described in our previous work (Zalesky et al. 2014). Briefly, we tested the top 20% of all 6,670 connections defined by the 116-region Automated Anatomical Labeling (AAL) atlas for evidence of time-varying connectivity (Tzourio-Mazoyer et al. 2002). A tapered sliding-window was adopted to enhance the suppression of spurious correlations and reduce sensitivity to outliers. The window length was set to 60 seconds and the exponent was set to a third of the window length (Leonardi and Van De Ville 2015; Pozzi et al. 2012; Zalesky and Breakspear 2015). To ensure that global efficiency values were not significantly different across brain parcellations, we performed the same analyses on a finer grained brain parcellation [200 volumetrically similar ROIs, top 20% of the 39,800 possible connections (Craddock et al. 2012)]. Efficiency values across time were highly correlated between the two brain parcellations.

### Searchlight decoding

For each participant, we implemented a spherical searchlight approach to decode between face and motion stimuli (Kriegeskorte et al. 2006). For each location within the whole brain mask, the voxel at the centre and its neighbors contained within the searchlight sphere (5 voxel diameter, 57 voxels volume) were extracted. The fMRI data from the resulting subset of voxels constituted the input to a linear support vector machine (Chang and Lin 2013). The fMRI data consisted of the 2 × *k* 3D volumes, where 2 is the number of fMRI sessions and *k* the total number of temporal subdivisions (i.e., 47). In each of the two fMRI sessions there were 48 trials (24 faces and 24 motion), each lasting for an average of 12 seconds. One trial was lost due to the use of the sliding window approach to measure dynamic functional connectivity, leaving a total of 47 trials in each fMRI session. Single-subject data from the second half of the first fMRI session and the first half of the second fMRI session were used to train the classifier (**Fig. 1**). Conversely, data from the first half of the first session and second half of the second session were used to test the classifier (**Fig. 1**). Single-subject maps consisted of values of decoding accuracy resulting from the cross-validation procedure (decimal values from 0 to 1, with 1 indicating 100% of correct labels). To evaluate decoding accuracy relative to chance (50% for the two stimulus categories of faces and motion), we employed permutation tests to construct a null distribution *for each participant*. A category-specific permutation was generated by shuffling the trial labels (i.e., face and motion). A whole brain, *single-subject* searchlight analysis was subsequently performed on the permuted data to create *single-subject chance accuracy maps*. This procedure, comprising permutation and searchlight analysis, was repeated 100 times per participant. Previous analyses have indicated that this number of repetitions is sufficient to achieve reliable estimation of false positive results (Stelzer et al. 2013).

To isolate brain regions containing significant information about the two stimulus categories at the group level (with one participant left out) in each leave-one-out run (Friedman et al. 2001), we adopted a validated non-parametric method (Stelzer et al. 2013). The voxel-wise average of the nonpermuted accuracy maps over the group formed an observed group level accuracy map. Then a bootstrapping method was used to construct a null distribution of the voxel-wise *group level* accuracy map from the 100 chance accuracy maps obtained previously. Each participant’s accuracy maps obtained from the permutations were recombined into *group-level* chance accuracy maps. This was achieved by randomly selecting one of the 100 chance accuracy maps for each participant. This selection was averaged to one permuted group-level accuracy map, and the bootstrapping procedure was repeated 10^5^ times. From the 10^5^ permuted maps arising from the bootstrapping procedure, a voxel-wise null distribution map of the *observed group-level accuracy* was built. For each voxel an accuracy threshold corresponding to p < 0.001 was then estimated to construct a voxel-wise observed significance threshold map. This initial p-value was established based on previous analyses (Stelzer etal. 2013). A voxel with an accuracy above the p < 0.001 threshold meant that it had better decoding performance than chance *at the group level*. The resulting map was used to binarise the observed group level accuracy map and the 10^5^ permuted maps. A cluster search was then performed on the observed and permuted binarised maps. In the cluster search algorithm two voxels were considered part of the same cluster if, and only if, they shared a face. We recorded all occurring cluster sizes in the observed and permuted maps. This procedure allowed us to estimate the probability of cluster size occurrence in the observed map. For each cluster size, a p-value was assigned and a significance threshold of cluster size was determined. Multiple comparison correction was implemented on all cluster p-values based on False Discovery Rate (FDR). A FDR-corrected cluster size threshold was then applied to the observed *group level* accuracy map. The resulting clusters formed an accuracy map comprising voxels with significant decoding at the group level (p < 0.05 FDR corrected). This map was used in subsequent region-based decoding for the left out participant.

### Effect of fluctuations in global efficiency on decoding

In each of the 20 leave-one-out runs, the significant group-level map defined using the searchlight approach (see above) was used as a feature set to calculate decoding accuracy as a function of changes in global network efficiency (for a schematic of the procedure, see SFig. 1). The BOLD signal of the left-out participant was used to both train (using the training epochs) and test (on the test epochs) a linear classifier. This procedure ensured that the selected regions of interest (ROI) within the thresholded map were significant at the group-level, and could be generalized to the whole cohort. Note, however, that the classifier was run within-participants (i.e., for each classifier, the training and the test sets were from the same participant). Decoding accuracy for a specific ROI in the left-out participant was calculated using the fMRI data corresponding separately to epochs of low global efficiency (lower 35%, 16 epochs) and high global efficiency (top 35%, 16 epochs) (SFig. 1). Significant differences in average ROI-based decoding performance between the low- and high-global efficiency states were tested for the 20 participants using a right-tailed nonparametric approach (based on the *a priori* prediction – recorded at the beginning of the study - that states of high global efficiency should increase neural decoding accuracy; see Introduction).

## Results

### Brain activity evoked by face and motion stimuli

Standard univariate analyses contrasting neural activity evoked by the two stimulus categories (face *minus* motion, and vice versa) showed that passive viewing of face stimuli evoked greater bloodoxygen- level dependent (BOLD) signal in bilateral fusiform gyrus [corresponding to the fusiform face area (Kanwisher et al. 1997)] than viewing of visual motion. Conversely, passive viewing of radial dot motion elicited greater BOLD responses in the middle temporal gyrus and the banks of the inferior temporal sulcus [corresponding to functional area MT (Annese et al. 2005; Walters et al. 2003)] than viewing of faces (p < 0.05 FDR corrected at cluster level; see **Fig. 2**). These results replicate what has been consistently observed in previous fMRI work assessing functionally specialised brain areas involved in the processing of human faces and visual motion (DeAngelis et al. 1998; Grill-Spector et al. 2006; Haxby et al. 2001; McGugin et al. 2012).

**Figure 2.**
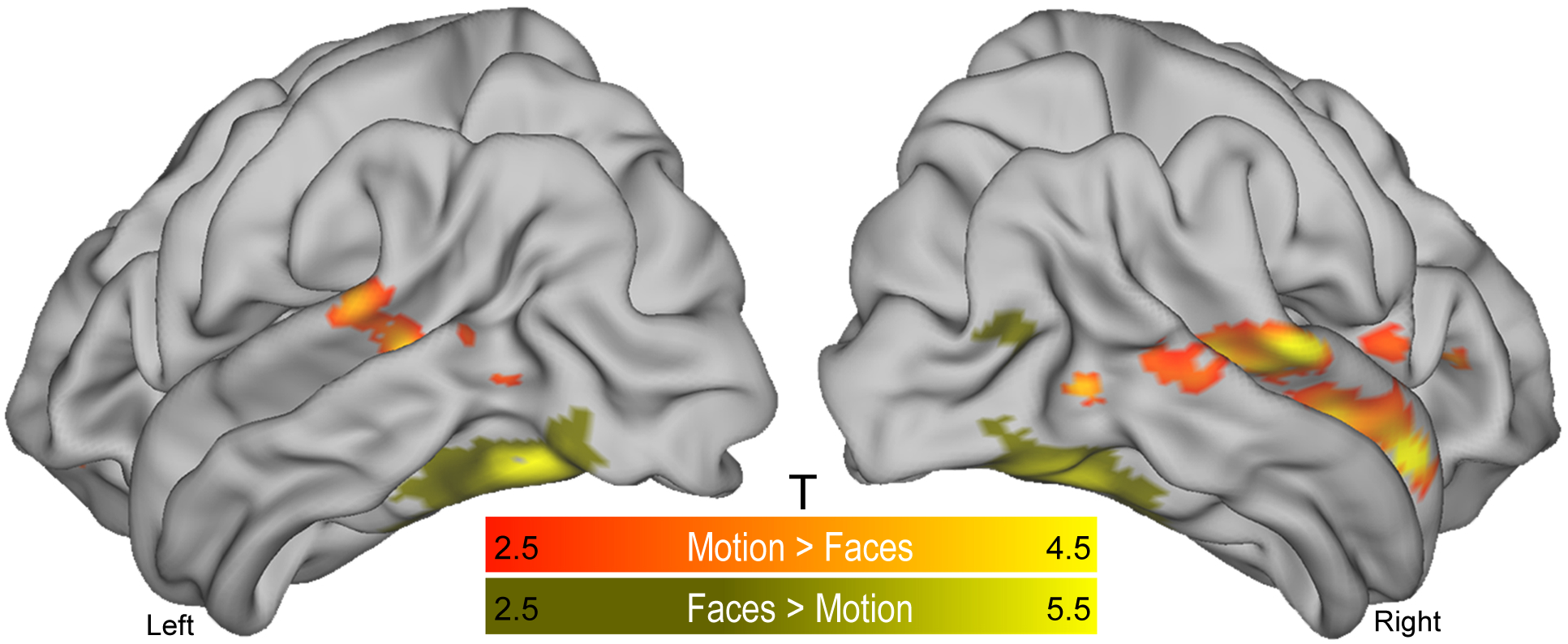
Univariate results. Brain regions showing significantly higher group-level (N = 20) activity during the presentation of face stimuli (yellow shading) than motion stimuli, and vice versa (orange shading) (second experimental session). Clusters were isolated using a standard general linear model framework (p < 0.05 False Discovery Rate corrected at the cluster level). As expected, the appearance of faces elicited greater activation in the fusiform gyrus bilaterally, whereas the motion stimuli elicited greater activation along the bank of the inferior temporal sulcus corresponding to human functional area MT (Annese et al. 2005; Walters et al. 2003). The appearance of motion stimuli was also associated with greater activation in regions comprising the middle temporal gyrus and frontal cortex. T = t-statistics.

### Decoding analyses of evoked responses to face and motion stimuli

Results of the decoding analyses revealed a widespread set of visual and non-visual brain regions with above-chance (> 53%) accuracy in distinguishing face and motion stimuli (**Fig. 3**). The bilateral fusiform gyri, area MT, and lateral portions of the inferior occipital cortex showed significant and very high (90-96%) decoding accuracy for the two categories of visual stimulus (p < 0.05 FDR corrected, **Fig. 3**). High (~ 80%) decoding accuracy was also observed in the occipital, inferior parietal, and posterior temporal cortices (**Fig. 3**). On the other hand, cortical clusters comprising prefrontal, frontal, mid-anterior temporal and superior parietal cortices displayed significant but relatively modest decoding accuracy (~ 55-70%, **Fig. 3**). These findings highlight the different degree of specialization in processing the two categories of visual stimuli, with sensory visual areas showing high specialization and associative regions showing a low degree of specialization.

**Figure 3.**
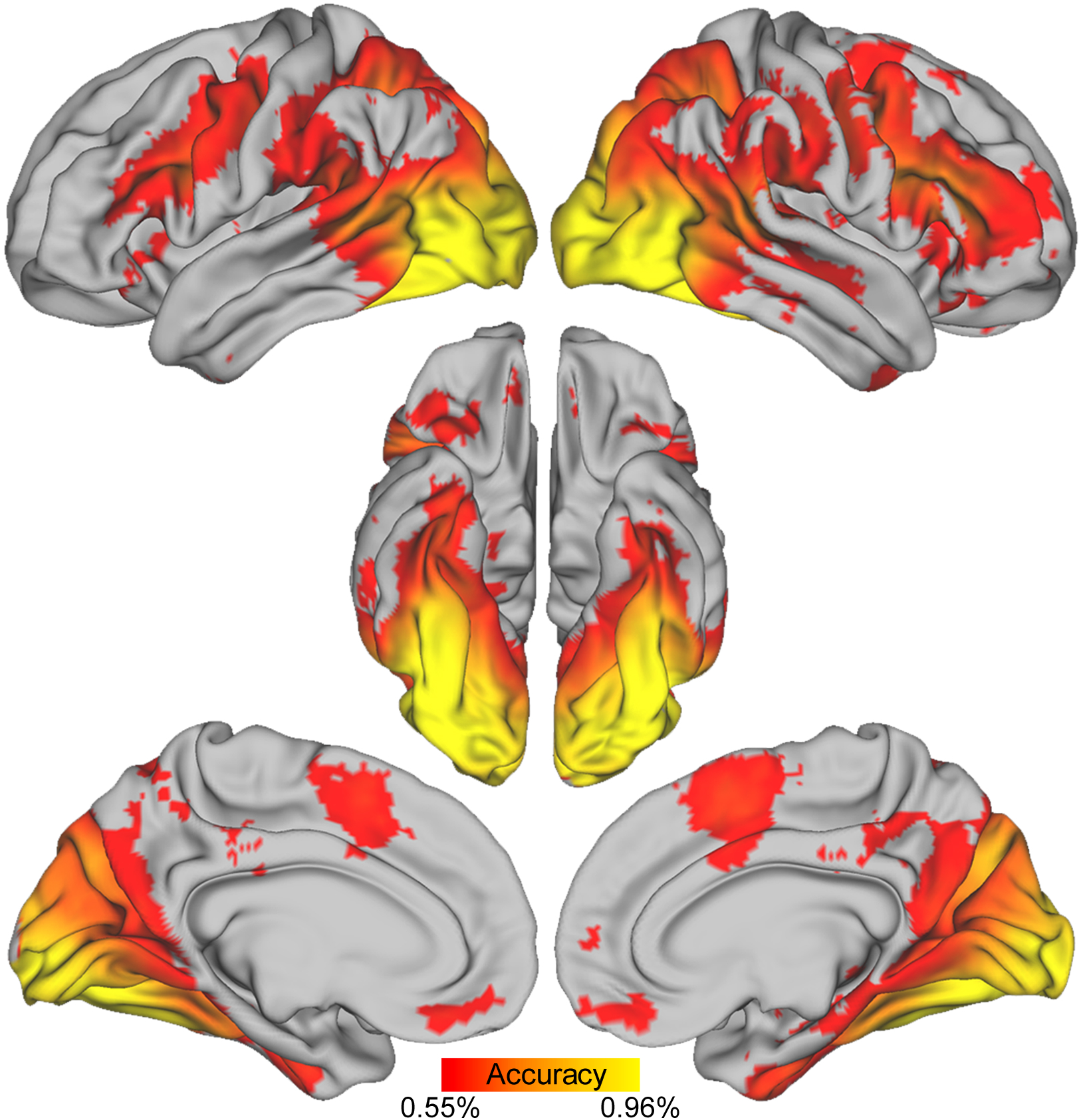
Multivariate results. Cortical regions showing reliable decoding between face and motionstimuli (group level map on the full sample). Decoding accuracy varied between 54% (> 54% depicted for illustration purposes) and 96%, as indicated by differences in red-yellow shading. Themap was generated using a standard searchlight decoding approach and a novel method allowing second level inferences (details in the Methods section).

### Time-resolved functional brain connectivity

Time-resolved functional connectivity was analysed using a sliding-window correlation approach. In line with recent methodological work (Leonardi and Van De Ville 2015; Zalesky and Breakspear 2015), we used a tapered 60-second window (86 time points per window; details in the Materials and Methods section). For each participant, this yielded a time series of functional connectivity estimates at a temporal resolution of 700 ms over a 20-min interval. Functional connectivity (**Fig. 4a**) and global efficiency (**Fig. 4b**) showed regular transitions between both highly and poorly connected neural states over time during the experimental session. As previously shown in a canonical restingstate context (Zalesky et al. 2014), our results revealed that spatially distributed brain regions transit in and out of states of global coordination over a timescale of several seconds. We previously showed that such changes in global neural states are not merely due to non-neural influences such as uncontrolled changes in blood pressure, respiration, heart rate or other physiological confounds (Zalesky et al. 2014). Furthermore, we found that dynamic fluctuations in functional connectivity obtained using two commonly used brain parcellations, the AAL (Tzourio-Mazoyer et al. 2002) and the “Craddock- 200” (Craddock et al. 2012), were highly correlated across participants and sessions (median r > 0.86, p < 0.001). Subsequent analyses were performed using network efficiency values derived from the AAL parcellation.

**Figure 4.**
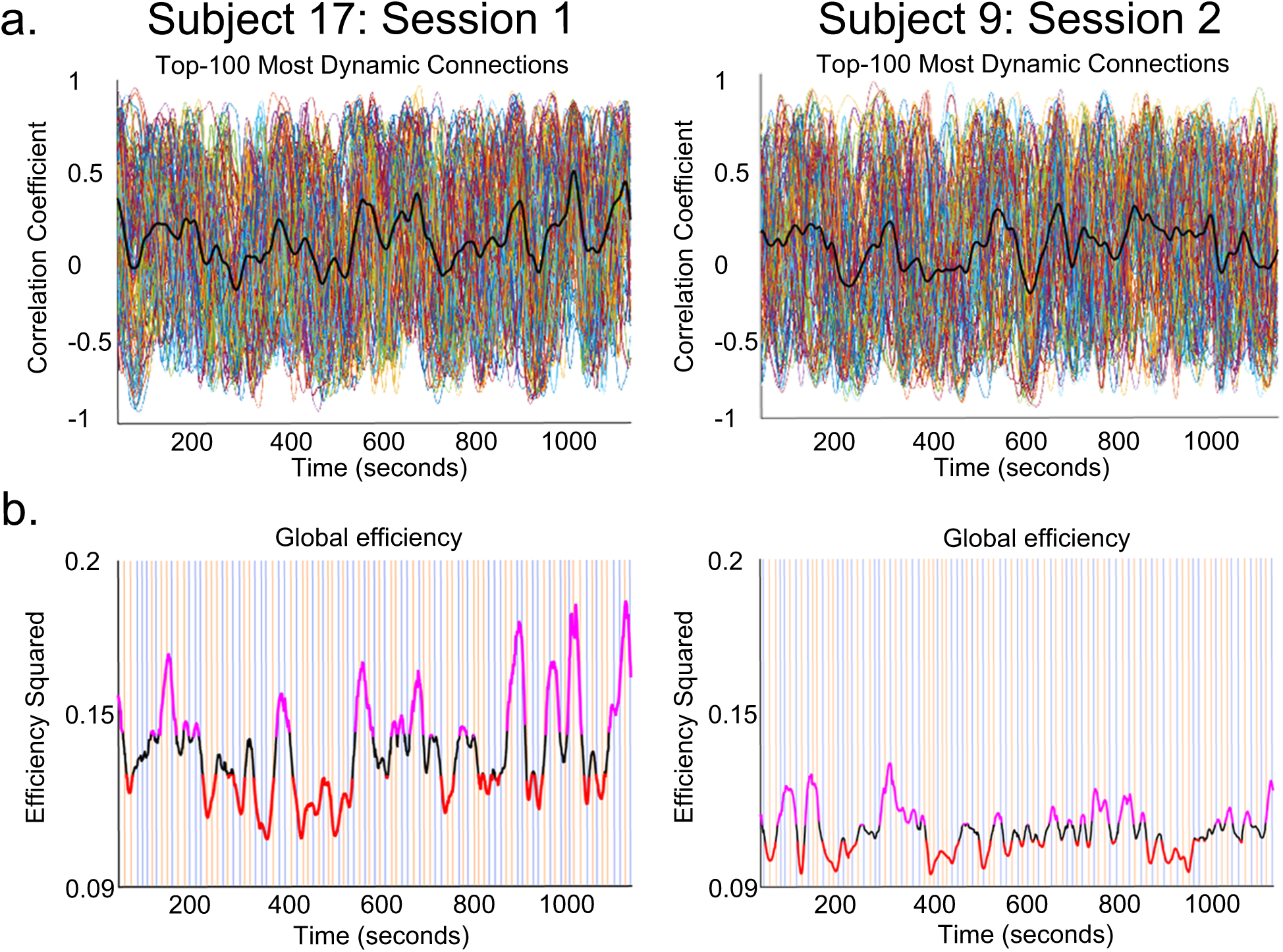
Dynamic functional connectivity. **(a)** Fluctuations in the time-series correlations of two representative participants, for a first and second fMRI session. Timeseries of correlation coefficients relating to the top-100 most dynamic connections (each colored line corresponds to a pairwise connection). Thick black lines represent mean functional connectivity. Functional connectivity across the two 20-minute fMRI sessions changes spontaneously, causing the emergence of global states of high and low network efficiency. **(b)** The orange and blue vertical lines indicate the presentation times of the two categories of visual stimuli (faces and motion). Stimuli were different within and between sessions. The thick black line represents fluctuations in global efficiency based on all pairwise connections (116 × 115= 13,340). Stimulus onset times were not correlated with peaks (top 35%, in purple; bottom 35%, in red) in global efficiency.

The two categories of visual stimulus (24 face and 24 motion stimuli in each fMRI session) were homogeneously distributed across the two fMRI sessions (see orange and blue lines in **Fig. 4b**). Stimulus onset times were not significantly correlated with peaks (volumes preceding a shift in the efficiency trend) in efficiency (**Fig. 4**) for any participants in the first or second fMRI sessions, except for one significant correlation out of 760. Likewise, the onsets of the highest (35%) and lowest (35%) states of global network efficiency (**Fig. 4**) were not correlated with the onsets of the stimuli. Thus, while the presented stimuli evoked the expected pattern of neural activity in functionally specialised brain clusters (**Fig. 2** and **Fig. 3**), stimulus onsets themselves did not influence slow fluctuations in global network activity.

### Impact of global network efficiency on stimulus decoding across the cortex

A significant difference in classifier performance was found between the high and low global efficiency states (p = 0.02 familywise error corrected, FWE; **Fig. 5a**). This performance difference in discriminating between faces and motion was identified in a map comprising all clusters showing significant decoding performance **(Fig. 3)**, excluding regions isolated by the mean subtraction analysis shown in **Fig. 2**. Specifically, the analysis showed that states of high whole-brain efficiency were associated with significantly better stimulus decoding performance than states of low global efficiency (**Fig. 5a and Fig. 5b**). Further analyses indicated that decoding accuracies within brain regions displaying different capacities to discriminate between face and motion events (i.e., 80 - 85% and 85 - 90% accuracy) across epochs were significantly correlated (r = 0.79, p = ~10^-10^). The absence of a difference in decoding performance in the clusters isolated by the univariate subtraction analysis was due to the very high accuracy of decoding in many of these brain regions (see **Fig. 2** and **Fig. 3** for decoding accuracy in the regions of interest) causing a ceiling effect in which the variance in performance was artificially limited. In fact, in these clusters the classifier’s accuracy was above 90% even in states of low global efficiency. These findings highlight the negligible impact of global fluctuations in functional connectivity on activity of highly specialised regions (such as the fusiform gyrus for face processing). On the other hand, less specialised brain regions evidently altered their activity patterns for the two stimulus categories as a function of changes in global network efficiency. Note that the variance in classifier performance was also very small in regions that showed close to chance decoding accuracy (53-55%, **Fig. 5b**). Removing these regions from the comparison map further increased the difference in decoding accuracy between states of high and low global efficiency.

**Figure 5.**
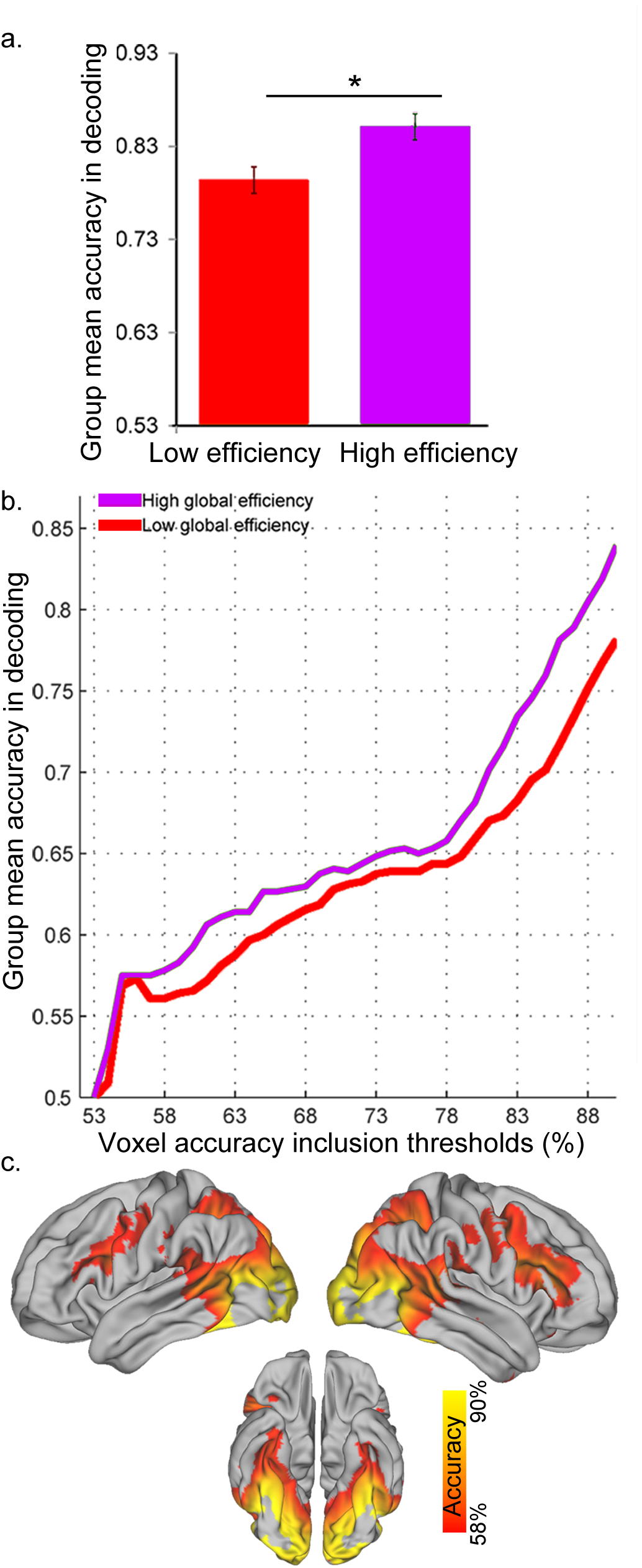
Decoding performance as a function of global efficiency. **(a)** Decoding accuracy between face and motion stimuli was significantly higher during epochs of high- versus low-global efficiency (* p = 0.02, familywise error corrected- FWE; error bars represent the SEM). **(b)** The observed main effect is based upon a consistent difference across accuracy thresholds, except for similar stimulus decoding accuracy at near-chance thresholds (53-55% accuracy). **(c)** Brain regions showing significant changes in decoding as a function of fluctuations in global brain efficiency. Note that regions with higher decoding accuracy were localised in the visual sensory areas, with fusiform gyrus and MT showing a decoding accuracy above 90% (not depicted) in both low and high network efficiency states.

To ensure that changes in global network efficiency were not simply related to a global change in the voxel-wise amplitude of the BOLD signal, we tested for a possible relationship between these two measures. There was *no* significant correlation between voxel-wise fluctuations in global efficiency and mean BOLD signal (p > 0.05). Furthermore, neither the mean BOLD signal across all grey matter voxels, nor the mean variance of the BOLD signal, differed significantly between high and low statesof global efficiency (t-tests, p > 0.05 with Bonferroni correction, with the exception of the variance of the BOLD signal for one participant, p = 0.0005). These results further support the notion that changes in global network efficiency (and decoding) are not due merely to fluctuations in the overall strength or variability of the BOLD signal (Zalesky et al. 2014).

Finally, we tested whether the observed difference in neural decoding as a function of global efficiency states could be explained by changes in non-neural signals. We examined decoding performance of head motion parameters (six values per time point) using a linear SVM and leave-one-out cross validation. Mean decoding accuracy across participants was close to chance for both global efficiency states (0.57 ±0.15 for the low efficiency state, and 0.52 ±0.15 for the high efficiency states), and there was no significant difference in mean decoding accuracy between them (t-test, p = 0.30). We also tested decoding performance of the white matter signal [values from 6321 voxels identified as white matter in the *ICBM-DTI-81* white-matter labels atlas (Hua et al. 2008)] using a linear SVM and leave-one-out cross validation. Mean decoding accuracy across participants was 0.38 ±0.14 for the low efficiency state and 0.35 ±0.13 for the high efficiency state. There was no significant difference in mean decoding accuracy between states (t-test, p = 0.60). These findings further support the notion that dynamic changes in global functional connectivity are driven by neural sources (Zalesky et al. 2014).

## Discussion

The brain is a complex, dynamic (non-stationary) network with rich spatiotemporal dynamics (Allen et al. 2014; Bassett et al. 2015; Gonzalez-Castillo et al. 2015; Shine et al. 2016; Zalesky et al. 2014). Disentangling interactions between neural dynamics occurring at different spatiotemporal scales remains a major endeavor for neuroscience. By combining time-resolved fMRI, network science, and a novel linear classification approach, we have shown that ultra-slow (< 0.15 Hz) time-varying fluctuations in global network efficiency modulate the extent to which local neural processes evoked by face and motion stimuli can be decoded from patterns of brain activity as measured with fMRI.

The functional link between ongoing global changes in brain connectivity and segregated decoding is reminiscent of findings from electroencephalography (EEG) studies, which suggest that global fluctuations in cortical synchrony preceding stimulus onset can facilitate local neural processes and visual perception. Typically, evidence for such facilitation has been found for transitions in brain rhythms comprising theta (Busch et al. 2009; Busch and VanRullen 2010; Landau and Fries 2012), alpha [(Mathewson et al. 2009; van Dijk et al. 2008), for a review see (Hanslmayr et al. 2011)], beta (Hipp et al. 2011), and gamma (Hipp et al. 2011) frequency bands. For example, participants’ perception of ambiguous audiovisual stimuli, and integration of auditory and visual information, is facilitated by enhanced beta and gamma band synchronization across widespread neural networks (Hipp et al. 2011). Our findings add to this emerging literature by suggesting that much slower spontaneous transitions in global neural synchronization can influence patterns of local neural activity elicited by two distinct categories of visual stimulus. This result opens the intriguing possibility that ultra-slow (< 0.15 Hz) global brain rhythms may play a causal role in the emergence of stimulus awareness in resting, and possibly active task, contexts, and is in line with recent work suggesting that global changes in fMRI connectivity relate to cognitive performance supported by segregated brain regions and systems (Shine et al. 2016).

Our analyses revealed that in addition to the fusiform gyrus (Haxby et al. 2001; Haxby et al. 1999) and area MT (Grossman and Blake 2002; Hong et al. 2012; Peelen et al. 2006; Seymour et al. 2009) awidespread set of visual and non-visual cortical regions is involved in discriminating human face and motion stimuli (**Fig. 3**). As such, the encoding of these distinct visual stimuli appears to rely on a diffuse neural network comprising regions at different levels of the cortical hierarchy. Specifically, our findings reveal that ongoing changes in global efficiency enhance the voxel-to-voxel mapping of stimulus-related information in brain regions comprising a diffuse set of visual and non-visual regions. The significant change in decoding performance across high- and low-efficiency states was not detected in category-specific regions such as the fusiform gyrus or area MT (**Fig. 2**), but instead was manifested in higher-level visual and non-visual regions such as the inferior parietal cortex and the frontal cortex (**Fig. 5c**). These findings suggest that the patterns of neural activity that distinguish face and motion stimuli are more evident across a widespread cortical network during states of high global efficiency (**Fig. 5c**), which might in turn be driven by enhanced communication between high-order cortical areas and low-level visual areas.

The sparse display of visual stimuli during scanning did not significantly influence global fluctuations in functional connectivity and efficiency. Moreover, the pseudorandom onsets of the two categories of visual stimulus were not related to fluctuations in pairwise functional connectivity and global efficiency. Likewise, in spite of an identical stimulus presentation sequence, dynamic fluctuations in connectivity were not correlated across fMRI sessions or participants. Thus, although slow fluctuations in network efficiency have a significant impact on local patterns of activity in both visual and non-visual areas, the opposite does not hold (i.e., the occasional appearance of visual events does not affect global network efficiency). Although this null effect of stimulus onset upon global network efficiency should be interpreted with caution, the apparent lack of influence of stimulus onsets on global efficiency fluctuations is consistent with recent computational work suggesting that global neural dynamics operating at ultra-slow timescales are relatively insensitive to faster dynamics in sensory nodes (Gollo et al. In press; Gollo et al. 2015; Kringelbach et al. 2015). Whether ultra-slow global fluctuations in connectivity isolated using fMRI are functionally independent or intermingled with faster, stimulus-evoked dynamics will require further investigation.

In this study we deliberately chose distinct categories of visual stimulus – faces and motion – to maximize the likelihood of uncovering significant differences in the strength of decoding across widespread regions of the cortex, given our sparse event-related design and relatively short resting scans. As a consequence, those cortical areas specialised for processing faces and motion – the fusiform region and area MT, respectively – showed ceiling-level decoding performance, as expected. In future work it will be important to determine whether the global changes in connectivity dynamics we report here also affect decoding accuracy for more subtle, within-category distinctions (e.g., between different facial expressions or motion directions). Based on the current results, it might be predicted that fluctuations in global brain connectivity should also modulate decoding accuracy for more subtle between-category distinctions, such as male versus female faces (Kaul et al. 2011) or different directions of visual motion (van Kemenade et al. 2014). Finally, while the current study was designed exclusively to assess physiological changes in local decoding across different states of global network efficiency, future work should determine whether participants’ behavioural performance (e.g., discrimination threshold) varies with fluctuations in network efficiency (Sadaghiani et al. 2015).

In summary, we have shown that decoding of neural activity elicited by two distinct categories of visual stimulus is facilitated during states of high– versus low-global efficiency. This boost in physiological changes linked to fMRI decoding was apparent in both visual and non-visual brain areas across several levels of the cortical hierarchy. Overall, our findings are in line with an emerging view that slow fluctuations in spontaneous neural activity underpinning global network efficiency play a critical role in modulating local patterns of neural activity while maintaining global network stability (Deco and Jirsa 2012; Deco et al. 2011; Gollo et al. In press; Gollo et al. 2015; Shine et al. 2016).

## Acknowledgments

The authors declare no competing financial interests. This study was supported by an ECR grant from The University of Queensland, Australia awarded to L. C. J. B. M was supported by an Australian Research Council (ARC) Australian Laureate Fellowship (FL110100103) and the ARC Centre of Excellence for Integrative Brain Function (ARC Centre Grant CE140100007). L. C., A. Z., and LLG were supported by the Australian National Health Medical Research Council (L. C., A. Z., J. B. M. APP1099082, A. Z. APP1047648, L. L. G. APP1110975). The authors would like to thank Dr Eve Dupierrix, Dr David Lloyd and Prof Michael Breakspear for helping generating the stimuli and commenting on earlier drafts of the manuscript. The authors also thank Siemens Healthcare for providing the prototype multiband sequence and Dr Kieran O’Brien for performing the sequence optimizations.

## Author Contributions

LC and JBM designed the research; LC and LH performed the data collection; LC, ZY, AZ, JS, and LH contributed to new analytic tools; LC and ZY analyzed the data. All authors wrote the paper.

## Supplementary Information

**SFigure 1.** *Analysis flowchart: testing of decoding accuracy for faces versus motion in states of high and low global network efficienc*y. Overview of the analysis pipeline.

**SMovie.** *Video presentation of a trial example.* Neutral human faces or radially moving dots sporadically and gradually appeared and disappeared. Participants were instructed to maintain their gaze on the white fixation cross, relax and ignore the stimuli. There was no task and no responses were required.

## References

Achard S, Bullmore E. 2007. Efficiency and Cost of Economical Brain Functional Networks. PLoS Comput Biol 3(2):e17.

Allen EA, Damaraju E, Plis SM, Erhardt EB, Eichele T, Calhoun VD. 2014. Tracking whole-brain connectivity dynamics in the resting state. Cereb Cortex 24(3):663–76.

Annese J, Gazzaniga MS, Toga AW. 2005. Localization of the human cortical visual area MT based on computer aided histological analysis. Cereb Cortex 15(7):1044–53.

Ashburner J. 2007. A fast diffeomorphic image registration algorithm. Neuroimage 38(1):95–113.

Bassett DS, Yang M, Wymbs NF, Grafton ST. 2015. Learning-induced autonomy of sensorimotor systems. Nat Neurosci 18(5):744–51.

Behzadi Y, Restom K, Liau J, Liu TT. 2007. A component based noise correction method (CompCor) for BOLD and perfusion based fMRI. Neuroimage 37(1):90–101.

Boubela RN, Kalcher K, Huf W, Kronnerwetter C, Filzmoser P, Moser E. 2013. Beyond Noise: Using Temporal ICA to Extract Meaningful Information from High-Frequency fMRI Signal Fluctuations during Rest. Front Hum Neurosci 7:168.

Breakspear M. 2004. "Dynamic" connectivity in neural systems: theoretical and empirical considerations. Neuroinformatics 2(2):205–26.

Bullmore E, Sporns O. 2012. The economy of brain network organization. Nat Rev Neurosci 13(5):336–49.

Busch NA, Dubois J, VanRullen R. 2009. The phase of ongoing EEG oscillations predicts visual perception. J Neurosci 29(24):7869–76.

Busch NA, VanRullen R. 2010. Spontaneous EEG oscillations reveal periodic sampling of visual attention. Proc Natl Acad Sci U S A 107(37):16048–53.

Chang C, Glover GH. 2010. Time-frequency dynamics of resting-state brain connectivity measured with fMRI. Neuroimage 50(1):81–98.

Chang CC, Lin CJ. LIBSVM: A Library for Support Vector Machines.

Chao-Gan Y, Yu-Feng Z. 2010. DPARSF: A MATLAB Toolbox for "Pipeline" Data Analysis of Resting-State fMRI. Front Syst Neurosci 4:13.

Craddock RC, James GA, Holtzheimer PE, 3rd, Hu XP, Mayberg HS. 2012. A whole brain fMRI atlas generated via spatially constrained spectral clustering. Hum Brain Mapp 33(8):1914–28.

de Pasquale F, Della Penna S, Sporns O, Romani GL, Corbetta M. 2015. A Dynamic Core Network and Global Efficiency in the Resting Human Brain. Cereb Cortex.

DeAngelis GC, Cumming BG, Newsome WT. 1998. Cortical area MT and the perception of stereoscopic depth. Nature 394(6694):677–80.

Deco G, Jirsa VK. 2012. Ongoing cortical activity at rest: criticality, multistability, and ghost attractors. The Journal of neuroscience 32(10):3366–3375.

Deco G, Jirsa VK, McIntosh AR. 2011. Emerging concepts for the dynamical organization of resting-state activity in the brain. Nature Reviews Neuroscience 12(1):43–56.

Ebner NC. 2008. Age of face matters: age-group differences in ratings of young and old faces. Behav Res Methods 40(1):130–6.

Friedman J, Hastie T, Tibshirani R. The elements of statistical learning [Internet]. Berlin: Springer series in statistics; 2001.

Friston KJ, Holmes AP, Worsley KJ, Poline J, Frith CD, Frackowiak RS. 1994. Statistical parametric maps in functional imaging: a general linear approach. Human brain mapping 2(4):189–210.

Fukunaga M, Horovitz SG, van Gelderen P, de Zwart JA, Jansma JM, Ikonomidou VN, Chu R, Deckers RH, Leopold DA, Duyn JH. 2006. Large-amplitude, spatially correlated fluctuations in BOLD fMRI signals during extended rest and early sleep stages. Magn Reson Imaging 24(8):979–92.

Gollo LL, Roberts JA, Cocchi L In press. Mapping how local perturbations influence systems-level brain dynamics. NeuroImage.

Gollo LL, Zalesky A, Hutchison RM, van den Heuvel M, Breakspear M. 2015. Dwelling quietly in the rich club: brain network determinants of slow cortical fluctuations. Philos Trans R Soc Lond B Biol Sci 370(1668).

Gonzalez-Castillo J, Hoy CW, Handwerker DA, Robinson ME, Buchanan LC, Saad ZS, Bandettini PA. 2015. Tracking ongoing cognition in individuals using brief, whole-brain functional connectivity patterns. Proc Natl Acad Sci U S A 112(28):8762–7.

Grill-Spector K, Sayres R, Ress D. 2006. High-resolution imaging reveals highly selective nonface clusters in the fusiform face area. Nat Neurosci 9(9):1177–85.

Grossman ED, Blake R. 2002. Brain Areas Active during Visual Perception of Biological Motion. Neuron 35(6):1167–75.

Hanslmayr S, Gross J, Klimesch W, Shapiro KL. 2011. The role of alpha oscillations in temporal attention. Brain Res Rev 67(1–2):331–43.

HaxbyJV, Gobbini MI, Furey ML, Ishai A, Schouten JL, Pietrini P. 2001. Distributed and overlapping representations of faces and objects in ventral temporal cortex. Science 293(5539):2425–30.

Haxby JV, Ungerleider LG, Clark VP, Schouten JL, Hoffman EA, Martin A. 1999. The effect of face inversion on activity in human neural systems for face and object perception. Neuron 22(1):189–99.

Hipp JF, Engel AK, Siegel M. 2011. Oscillatory synchronization in large-scale cortical networks predicts perception. Neuron 69(2):387–96.

Hong SW, Tong F, Seiffert AE. 2012. Direction-selective patterns of activity in human visual cortex suggest common neural substrates for different types of motion. Neuropsychologia 50(4):514–21.

Hua K, Zhang JY, Wakana S, Jiang HY, Li X, Reich DS, Calabresi PA, Pekar JJ, van Zijl PCM, Mori S. 2008. Tract probability maps in stereotaxic spaces: Analyses of white matter anatomy and tract-specific quantification. Neuroimage 39(1):336–347.

Kanwisher N, McDermott J, Chun MM. 1997. The fusiform face area: a module in human extrastriate cortex specialized for face perception. J Neurosci 17(11):4302–11.

Karahanoglu FI, Van De, Ville D. 2015. Transient brain activity disentangles fMRI resting-state dynamics in terms of spatially and temporally overlapping networks. Nat Commun 6:7751.

Kaul C, Rees G, Ishai A. 2011. The gender of face stimuli is represented in multiple regions in the human brain. Frontiers in Human Neuroscience 4:238.

Kriegeskorte N, Goebel R, Bandettini P. 2006. Information-based functional brain mapping. Proc Natl Acad Sci U S A 103(10):3863–8.

Kringelbach ML, McIntosh AR, Ritter P, Jirsa VK, Deco G. 2015. The Rediscovery of Slowness: Exploring the Timing of Cognition. Trends Cogn Sci 19(10):616–28.

Landau AN, Fries P. 2012. Attention samples stimuli rhythmically. Curr Biol 22(11):1000–4.

Latora V, Marchiori M. 2001. Efficient behavior of small-world networks. Phys Rev Lett 87(19):198701.

Leonardi N, Van De, Ville D. 2015. On spurious and real fluctuations of dynamic functional connectivity during rest. Neuroimage 104:430–6.

Mathewson KE, Gratton G, Fabiani M, Beck DM, Ro T. 2009. To see or not to see: prestimulus alpha phase predicts visual awareness. J Neurosci 29(9):2725–32.

McGugin RW, Gatenby JC, Gore JC, Gauthier I. 2012. High-resolution imaging of expertise reveals reliable object selectivity in the fusiform face area related to perceptual performance. Proc Natl Acad Sci U S A 109(42):17063–8.

Murray JD, Bernacchia A, Freedman DJ, Romo R, Wallis JD, Cai X, Padoa-Schioppa C, Pasternak T, Seo H, Lee D et al. 2014. A hierarchy of intrinsic timescales across primate cortex. Nat Neurosci 17(12):1661–3.

Niazy RK, Xie J, Miller K, Beckmann CF, Smith SM. 2011. Spectral characteristics of resting state networks. Prog Brain Res 193:259–76.

Peelen MV, Wiggett AJ, Downing PE. 2006. Patterns of fMRI activity dissociate overlapping functional brain areas that respond to biological motion. Neuron 49(6):815–22.

Ponce-Alvarez A, Deco G, Hagmann P, Romani GL, Mantini D, Corbetta M. 2015. Resting-state temporal synchronization networks emerge from connectivity topology and heterogeneity. PLoS Comput Biol 11(2):e1004100.

Power JD, Barnes KA, Snyder AZ, Schlaggar BL, Petersen SE. 2012. Spurious but systematic correlations in functional connectivity MRI networks arise from subject motion. Neuroimage 59(3):2142–54.

Power JD, Mitra A, Laumann TO, Snyder AZ, Schlaggar BL, Petersen SE. 2014. Methods to detect, characterize, and remove motion artifact in resting state fMRI. Neuroimage 84:320–41.

Pozzi F, Di Matteo T, Aste T. 2012. Exponential smoothing weighted correlations (vol 85, 175, 2012). European Physical Journal B 85(8).

Sadaghiani S, Poline JB, Kleinschmidt A, D'Esposito M. 2015. Ongoing dynamics in large-scale functional connectivity predict perception. Proc Natl Acad Sci U S A 112(27):8463–8.

Seymour K, Clifford CW, Logothetis NK, Bartels A. 2009. The coding of color, motion, and their conjunction in the human visual cortex. Current Biology 19(3):177–183.

Shine JM, Bissett PG, Bell PT, Koyejo O, Balsters JH, Gorgolewski KJ, Moodie CA, Poldrack RA. 2016. The Dynamics of Functional Brain Networks: Integrated Network States during Cognitive Task Performance. Neuron.

Sporns O. 2011. Networks of the brain. Cambridge: Massachusetts Institute of Technology.

Stelzer J, Chen Y, Turner R. 2013. Statistical inference and multiple testing correction in classification-based multi-voxel pattern analysis (MVPA): random permutations and cluster size control. Neuroimage 65:69–82.

Sun FT, Miller LM, D'Esposito M. 2004. Measuring interregional functional connectivity using coherence and partial coherence analyses of fMRI data. Neuroimage 21(2):647–58.

Tagliazucchi E, Laufs H. 2014. Decoding wakefulness levels from typical fMRI resting-state data reveals reliable drifts between wakefulness and sleep. Neuron 82(3):695–708.

Tzourio-Mazoyer N, Landeau B, Papathanassiou D, Crivello F, Etard O, Delcroix N, Mazoyer B, Joliot M. 2002. Automated anatomical labeling of activations in SPM using a macroscopic anatomical parcellation of the MNI MRI single-subject brain. Neuroimage 15(1):273–89.

van Dijk H, Schoffelen JM, Oostenveld R, Jensen O. 2008. Prestimulus oscillatory activity in the alpha band predicts visual discrimination ability. J Neurosci 28(8):1816–23.

van Kemenade BM, Seymour K, Christophel TB, Rothkirch M, Sterzer P. 2014. Decoding pattern motion information in V1. cortex 57:177–187.

Walters NB, Egan GF, Kril JJ, Kean M, Waley P, Jenkinson M, Watson JD. 2003. In vivo identification of human cortical areas using high-resolution MRI: an approach to cerebral structure-function correlation. Proc Natl Acad Sci U S A 100(5):2981–6.

Zalesky A, Breakspear M. 2015. Towards a statistical test for functional connectivity dynamics. Neuroimage 114:466–70.

Zalesky A, Fornito A, Cocchi L, Gollo LL, Breakspear M. 2014. Time-resolved resting-state brain networks. Proc Natl Acad Sci U S A 111(28):10341–6.

